# A three year follow-up study of gadolinium enhanced and non-enhanced regions in multiple sclerosis lesions using a multi-compartment *T*_2_ relaxometry model

**DOI:** 10.1101/365379

**Authors:** Sudhanya Chatterjee, Olivier Commowick, Onur Afacan, Benoit Combes, Simon K. Warfield, Christian Barillot

## Abstract

Demyelination, axonal damage and inflammation are critical indicators of the onset and progress of neurodegenerative diseases such as multiple sclerosis (MS) in patients. Due to physical limitations of imaging such as acquisition time and imaging resolution, a voxel in a MR image is heterogeneous in terms of tissue microstructure such as myelin, axons, intra and extra cellular fluids and free water. We present a multi-compartment tissue model which estimates the water fraction (WF) of tissues with short, medium and high *T*_2_ relaxation times in a *T*_2_ relaxometry MRI voxel. The proposed method is validated on test-retest data of healthy controls. This model was then used to study longitudinal trends of the tissue microstructures for two sub-regions of the lesions: gadolinium enhanced (*E*+) and non-enhanced (*L*–) regions of MS lesions in 10 MS patients over a period of three years. The water fraction values in *E*+ and *L*– regions were found to be significantly different (*p* < 0.05) over the period of first three months. The results of this study also showed that the estimates of the proposed *T*_2_ relaxometry model on brain tissue microstructures have potential to distinguish between regions undergoing active blood brain barrier breakdown from the other regions of the lesion.

## 1 Introduction

Magnetic resonance imaging (MRI) is one of the most widely used in-vivo imaging method for obtaining information on brain health. However, MRI voxels have limited resolution due to physical constraints. Use of advanced MRI techniques such as *T*_2_ relaxometry and diffusion weighted MRI can address this issue by providing quantitative estimates on brain tissue microstructures such as myelinated and unmyelinated axons, brain fiber orientation etc [1]. In patients with neurodegenerative diseases (such as multiple sclerosis (MS)), obtaining information on health of axons and myelin using in-vivo imaging techniques can improve our understanding of the current status and the progress of the disease in the patients. Obtaining this information can provide critical knowledge on the brain health.

The myelin, axons and fluids in the brain can be distinguished based on the nature of their water content. Myelin is a tightly wrapped structure around the axons. The water in myelin is very tightly bound to its surface. Hence myelin has a very short *T*_2_ relaxation time compared to the other brain tissue structures. The axons, gray matter cells and other cells in the brain have a higher *T*_2_ relaxation time than that of myelin but less than that of the fluids. Free fluids have the largest *T*_2_ relaxation time. Hence, based on the *T*_2_ relaxation time brain tissues can be broadly classified into three categories, short-, medium– and high–*T*_2_ components [2,3]. Since a voxel in a MR image of the brain is heterogeneous, there is a certain amount of the three *T*_2_ components present in each voxel. In addition to these, in presence of an infection there might also be some fluid accumulation. Hence a quantitative metric that conveys information on the condition of these tissues can provide useful insights into the current brain tissue health.

Quantitative *T*_2_ relaxometry MRI sequences offer the capability to distinguish tissues based on their *T*_2_ relaxation times. *T*_2_ relaxometry MRI has been used effectively to calculate the myelin water fraction using a variety of approaches. Whittall et al. and MacKay et al. [2, 4] obtained myelin water fraction using a multi-component *T*_2_ model [5]. The multicomponent *T*_2_ model describes the observed *T*_2_ decay curve as a weighted sum of an arbitrary number of decay curves. The weight of each *T*_2_ decay curve is obtained using non-negative least squares (NNLS). Whittall et al. obtained myelin water fraction of the brain tissues by assuming the *T*_2_ values of water in myelin to be in the range of 10 – 55 milliseconds [2]. The multi-component *T*_2_ models select an arbitrary number of *T*_2_ decay curves to model the observed decay signal [5]. For example, Whittall et al. [2] considered 80 logarithmically spaced values between 15 milliseconds and 2 seconds. Since the number of parameters to be estimated considerably outweighs the number of observations (i.e. number of echoes), some regularization is mandatory. However, the choice of the regularization term and its extent affects the fitting measure [6].

An alternate approach is a multi-compartment *T*_2_ relaxometry model where the different *T*_2_ water pools in brain tissues are modeled as a weighted mixture of pre-defined continuous functions. Stanisz and Henkelman [7] fitted two *T*_2_ components (short and long *T*_2_ in bovine optic nerve), each having a gaussian distribution on a logarithmic scale. Akhondi-Asl et al. proposed a framework to calculate myelin water fraction by modeling the R_2_ space as a weighted mixture of inverse Gaussian distribution of fast–, medium– and slow– decaying components with respect to *T*_2_ relaxation times [8]. Layton et al. proposed a maximum likelihood estimation approach to estimate the myelin water fraction values [9]. They obtained the Cramer-Rao lower bounds of the variables to establish the difficult nature of simultaneously estimating the model parameters and weights for each compartment for such models. In contrast to the multiple exponential *T*_2_ fitting approach, these models do not suffer from the ill-posed weights estimation problem. In these models, the regularization is performed a priori rather than a posteriori.

In this work, we propose a method for computing water fractions corresponding to fast–, medium– and slow– decaying components with respect to *T*_2_ relaxation times from *T*_2_ relaxometry MRI data acquired using 2D multislice Carr-Purcell-Meiboom-Gill (CPMG) sequence. In this estimation framework, the *T*_2_ space is modeled as a weighted mixture of three continuous probability density functions (PDF) representing the three *T*_2_ compartments. For such models, the robustness and accuracy of the implementations to simultaneously estimate the weights and parameters of the distributions was found to be non-trivial and not reliable [9]. We therefore chose to take advantage of the earlier studies performed in this field to fix the parameters of the distributions representing the *T*_2_ relaxation components [3, 10, 11]. Hence only the estimation of weights remain, making the estimation model robust. In our work, we need to correct for the effect of the stimulated echoes due to imperfect refocusing (due to the *B*_1_ inhomogeneities) that leads to errors in *T*_2_ estimation [12]. The EPG algorithm is used to account for these stimulated echoes [13]. The field inhomogeneity (*B*_1_) is estimated numerically. Since the *T*_2_ space is modeled as a weighted mixture of three continuous PDFs representing the three components, the proposed model does not include any regularization on the water fractions. The estimated weights of each compartment provide a quantitative estimate of the tissue microstructure in a voxel. Experiments were carried out to evaluate repeatability of the quantitative markers on healthy controls.

In the last part of this work, we show an application of the method on MS patients. We report observations on the evolution of the quantitative markers obtained from the model in enhancing (MS lesion regions undergoing active blood brain barrier breakdown) and non-enhancing MS lesion regions. Vavasour et al. studied the MWF and total water content in three new MS lesions in two patients over a year [14]. Levesque et al. studied evolution of MWF and geometric mean of *T*_2_ values of gadolinium enhancing lesions in five MS patients over a period of one year [15]. Although MWF was found to be quite informative in suggesting changes in active lesions, the geometric mean *T*_2_ values was sensitive to changes in the active lesions. Vargas et al. studied MWF of gadolinium enhancing and non-enhancing lesions [16]. Measurements were compared for two acquisitions with median duration of around 6 months between the scans. Authors found that contrary to the non-enhancing lesions, there was significant improvement in MWF of enhancing lesions between the two scans. Vargas et al. also mention that MWF is a combined measure of edema and demyelination [16]. The MWF is a relative measure and a change in its values should be studied in conjunction with the remaining water fraction measures. In our analysis, we observed and compared the change in water fractions of tissues with short, medium and high *T*_2_ values in enhancing and non-enhancing regions of lesions in 10 MS patients over a period of 3 years.

## 2 Materials and methods

### 2.1 Estimation framework

#### Signal model

We model the *T*_2_ space as a weighted mixture of three continuous probability density functions (PDF) representing the three *T*_2_ relaxometry compartments. The compartments represent tissues with short, medium and high *T*_2_ relaxation times. The weight of the *j*–th distribution is denoted by *w*_*j*_. The weights are normalized such that ∑_*j*_ *w*_*j*_ = 1.The signal of a voxel at the i–th echo time (*t*_*j*_) is therefore given as:

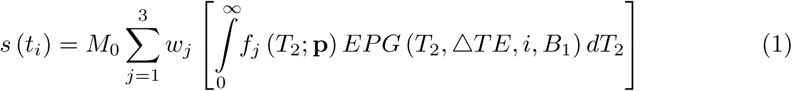

where *t*_*i*_ = *i* × Δ*TE* and Δ*TE* is the echo spacing. Each *f*_*j*_ (*T*_2_; **p**) is the chosen PDF with parameters **p**. As mentioned in the introduction, for robustness reasons, the PDF parameters are pre-selected and fixed keeping in mind histology findings reported in the literature [3, 11]. In Eq. (1), M_0_ is the magnetization constant. *EPG*(·) represents the stimulated echo computed at the time point (*t*_*i*_ = *i* × Δ*TE*) using the EPG algorithm [13]. *B*_1_ is the field inhomogeneity.

#### Optimization

Without any loss of generality, *M*_0_ and 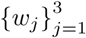 in Eq. (1) can be combined into a single term, 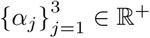. The weight corresponding to each compartment can then be obtained as, *w*_*j*_ = *a*_*j*_/∑_*i*_ *α*_*i*_ and M_0_ as *∑*_*i*_ *α*_*i*_. The signal of the voxel at time *t*_*i*_ is expressed as:

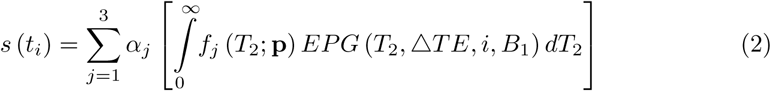

The parameters to be estimated in Eq. (2) are: 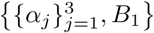. The optimization is thus formulated as a least square problem:

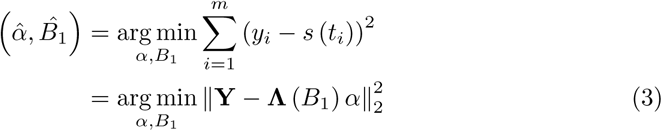

where *m* is the number of echoes; **Y** ∈ ℝ^*m*^ is the observed signal; *α* = {*α*_1_, *α*_2_, *α*_3_} ∈ ℝ^+^3^^. Each element of **⋀**

(⋀_*ij*_ = *λ*_*j*_ (*t*_*j*_; *B*_1_); *i* = {1,…, *m*}, *j* = {1, 2, 3}) in Eq. (3) is computed as:

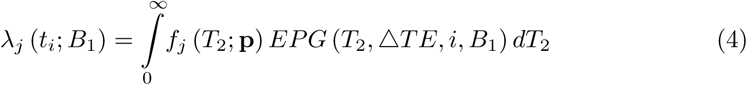

The parameters to be estimated in the least squares optimization problem stated in Eq. (3), a and *B*_1_, are linear and non-linear in nature respectively. However they are linearly separable. Hence optimization for a and *B*_1_ is performed alternatively until convergence is obtained in a desired error limit. In the first step, a is computed by non-negative least squares (NNLS) optimization [17] with a fixed *B*_1_ value. In the next step, the weights computed in the first step are used to compute *B*_1_ by a gradient free optimizer (BOBYQA). We choose to perform a numerical optimization to obtain *B*_1_ as it does not have any closed form solution [13]. The integral in Eq. (4) also does not have a closed form solution. Hence the integral is computed using the Riemann sum approach by dividing the *T*_2_ region into rectangles of finite width (= 0.33ms in our case) over the range of [0, 2500] ms.

In this work, the PDFs to represent each *T*_2_ relaxometry compartment 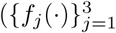 in Eq. (1)) are chosen as Gaussian PDFs. The mean and standard deviations are chosen for the three compartments based on histology findings reported in the literature [3, 10, 11] and are set as *μ* = {20,100, 2000} and a = {5,10, 80} (all values in milliseconds). The proposed method was implemented in C++ and is made available as an open source library `Anima`^1^

### 2.2 Experimental Methods

Repeatability is an important aspect of quantitative MRI techniques. In the first experiment we carried out test-retest experiments to assess the repeatability of the proposed method. An important motivation of this study is to gain more insights into the neurodegenerative disease phenomenon using the tissue microstructure information. Hence, in the second experiment we performed a 3-year follow-up study on 10 multiple sclerosis patients.

#### 2.2.1 Experiment-1: Repeatability test

The objective of this experiment is to observe whether the proposed model is repeatable in terms of estimation of the microstructure maps. For that purpose, test retest *T*_2_ relaxometry scans of 4 healthy controls were obtained to evaluate the repeatability of the proposed method.

##### Healthy controls

The age of the healthy controls was in the range of 26-32 years. 15 regions of interest (ROI) were marked in the brain for each healthy control over which the test and retest values of the compartments’ water fractions were compared. An illustration of these ROI on a subject is shown in Fig. 1.

**Fig 1.**
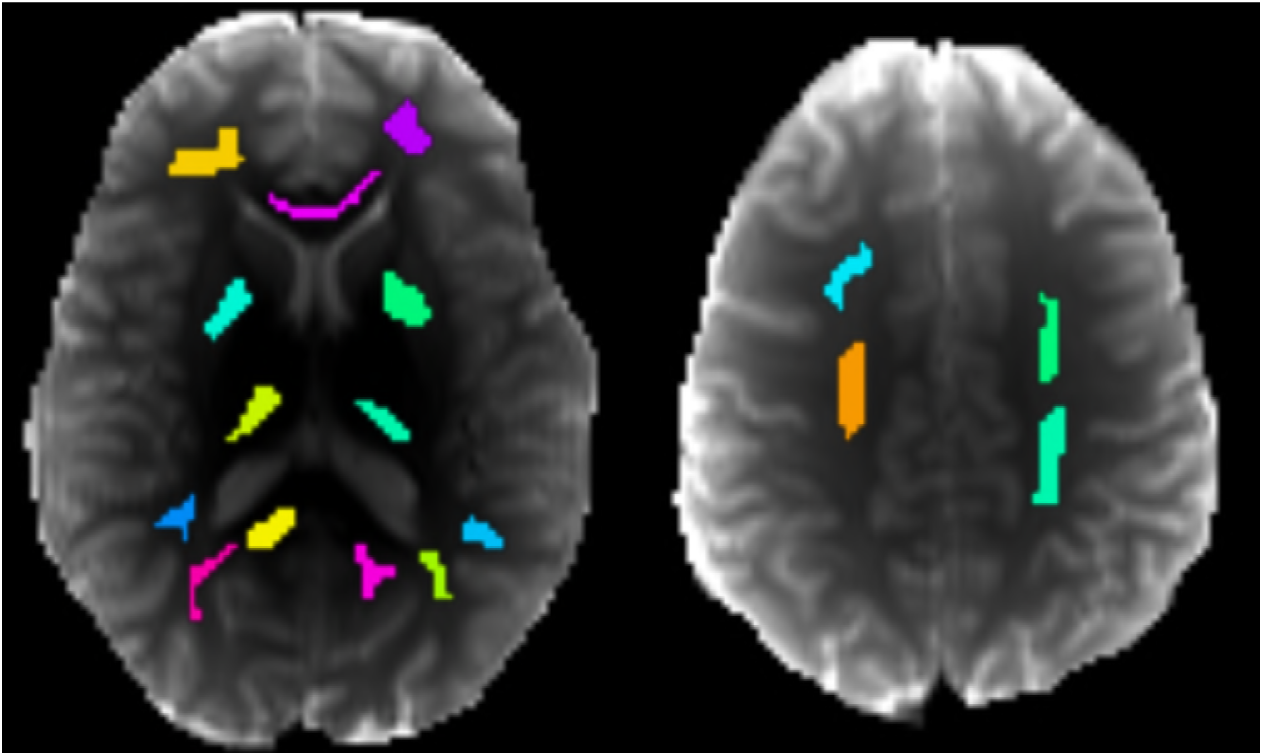
Assessing repeatability from test-retest data. Test retest scans were performed for 4 healthy controls to study the repeatability of the proposed method. This figure shows the 15 regions which were marked on the healthy controls over which the repeatability was studied.

##### Acquisition details

The details of the acquired data are as follows: 3T MRI scanner; 2D multislice CPMG sequence; 32 echoes; first echo at 9ms; Δ*TE* = 9ms; *TR* = 3000ms; single slice acquisition; in plane resolution= 1.1mm × 1.1mm; slice thickness of 4mm; matrix size of 192 × 192; number of averages= 1.

##### Analysis

A Bland-Altman plot was used to observe the repeatability of the estimations over the ROIs. From the plot we obtained the mean deviation (*m*_*d*_) of the test retest values for the ROIs and checked whether there are noticeable systematic changes in the estimations. We further look at the limits of agreement (LoA) between the test retest estimations from the plot.

#### 2.2.2 Experiment-2: Application to Multiple Sclerosis (MS)

Multiple sclerosis lesions are focal lesions and grow in a concentric manner [18]. In the early stages, brain tissues in the MS lesions undergo active blood brain barrier breakdown [18, 19]. Surrounding the core of the lesion is the edema as a result of tissue inflammation due to ongoing tissue damage. As compared to the normal appearing brain matter, the entire MS lesion regions appear hyper-intense on *T*_2_ weighted MRI. However, only the regions of the lesion undergoing active blood brain barrier breakdown appear hyperintense on *T*_1_ weighted MRI acquired post Gd injection. Hence, lesions in active state have two regions, a region which actively undergoes blood brain barrier breakdown and the regions which do not.

The objective of this experiment is to study the evolution of compartments’ water fractions in regions of lesions in MS patients undergoing active blood brain barrier breakdown and the regions which are not. In addition to that, we shall observe whether the water fraction values of the compartments for the two regions are in confirmation with the pathological knowledge of MS lesions.

##### Patient Data

We studied 10 MS patients over a period of 36 months. All patients included in this study had an episode of clinically isolated syndrome (CIS). The patient cohort comprised of an equal number of male and female subjects and their median age was 28 years. All participants of this study gave their written consent. Data was acquired at eight time points over a period of 36 months. An acquisition at the baseline was followed by acquisitions at 3, 6, 9, 12, 18, 24 and 36 months from the baseline. The lesions in patient scans were marked by an expert radiologist on *T*_2_ weighted images at the baseline. All participants were informed and provided their written consent for the study.

##### Acquisition details

All data was acquired on a 3T MRI scanner. The acquisition details for the *T*_2_ relaxometry data are as follows: 2D multislice CPMG sequence; 7 echoes; first echo at 13.8ms; Δ*TE* = 13.8ms; *TR* = 4530ms; voxel dimensions = 1.3 × 1.3 × 3.0 mm^3^; spacing between slices of 3mm; matrix size of 192 × 192; number of averages = 1; acquisition time ≈ 7 minutes. A *T*_1_ weighted scan was obtained post gadolinium injection to identify the lesions undergoing active blood brain barrier breakdown. The post Gadolinium injection (0.1mmol/kg gadopentate dimeglumine) acquisition details are as follows: transverse spin echo *T*_1_ weighted images; voxel dimensions = 1.0 × 1.0 × 3.0 mm^3^; spacing between slices of 3mm; matrix size of 256 × 256; number of averages = 1. The protocols were approved by the institutional review board of Rennes University Hospital.

##### Analysis

In this study, we define two groups of MS lesions, (a) *E*+: lesion regions which appear hyper-intense on post-Gd injection *T*_1_-weighted images and (b) *L*–: lesion regions appearing hyper-intense on *T*_2_-weighted images only. Thus a lesion might have both *E*+ and *L*– regions in it. An illustration of the lesion regions is shown in Fig. 2.

**Fig 2.**
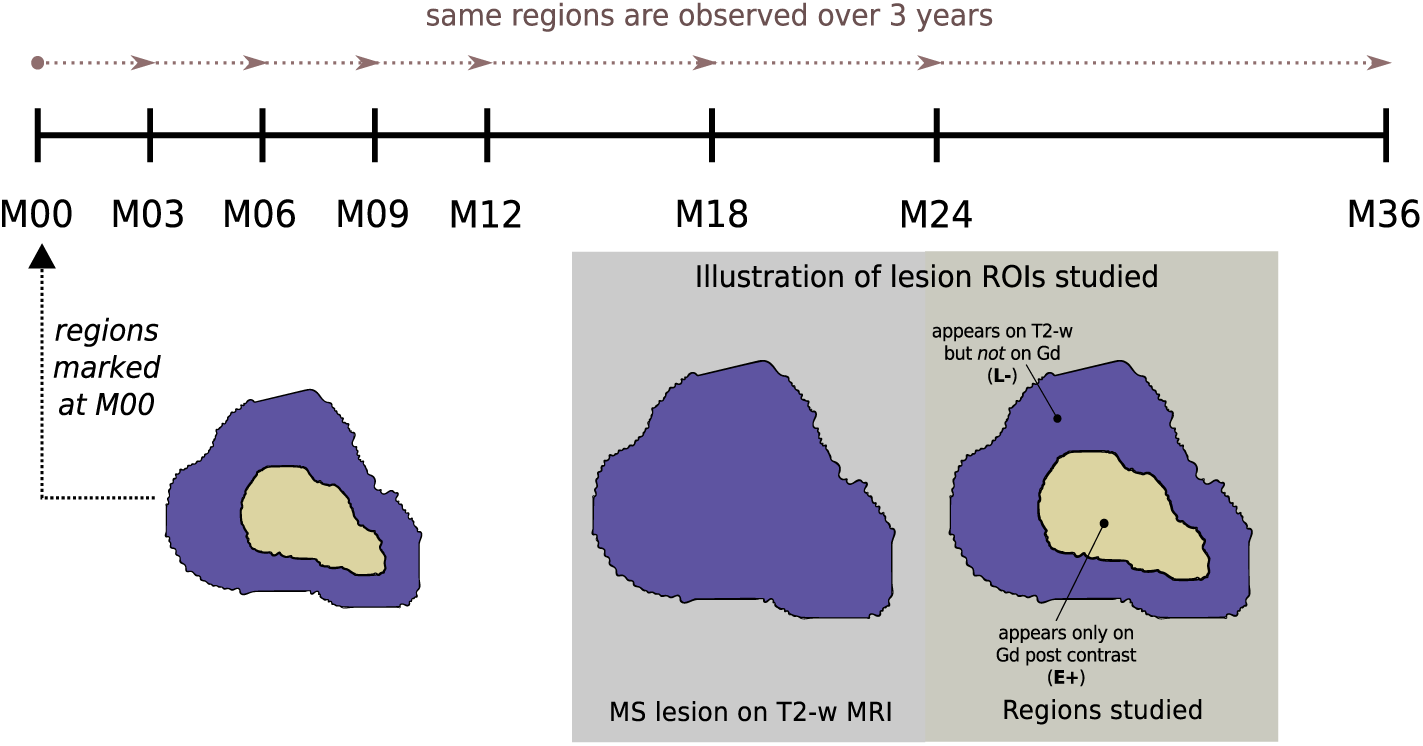
MS patient study design. The MS lesions were marked on *T*_2_ weighted on gadolinium post contrast *T*_1_ spin echo MR images of the patients acquired at the first visit (i.e. M00). Eight scans are obtained from time of the first visit over a period of 36 months for each patient at intervals shown in the figure. The region of interest (ROI) marked at M00 is studied over the period of 36 months. Two ROIs are studied in this work: (*i*) region of the lesion which appears on the gadolinium post contrast *T*_1_ weighted spin echo images. These are the regions of the lesions undergoing active blood brain barrier breakdown, (*ii*) lesion region appearing on *T*_2_ weighted MR images but not on gadolinium post contrast images.

The lesion ROIs are marked at baseline and the same region is observed over a period of three years. In the 10 MS patients, we observed 229 *L*– and 25 *E*+ lesion regions. Since the lesions were marked on *T*_2_ weighted images, all processed images were registered to the *T*_2_ images using a block matching algorithm [20, 21].

## 3 Results

### 3.1 Experiment-1: Repeatability test

The Bland-Altman (BA) plots for short, medium and high-*T*_2_ water fraction estimates over the ROIs are shown in Fig. 3. BA plots are scatter plots between the average test-retest measurement and the difference between test-retest measurements. We measured the means of the estimated values of 15 ROIs in 4 healthy controls. The BA plots for short, medium and high-*T*_2_ water fraction estimates for the 15 ROIs are shown in Fig. 3(a), 3(b) and 3(c) respectively. The plots shows the level of mean error (*m*_*d*_) observed. The gray area around the mean error level is its 95% confidence interval (CI). Along with *m*_*d*_, the *m*_*d*_ ± 1.96 × *σ*_d_ levels are also shown and are referred to as levels of agreement (LoA). *σ*_*d*_ is the standard deviation associated with the errors observed in the test-retest measurements. LoA is thus an empirical estimate of the range around *m*_*d*_ within which 95% of the differences are expected to exist. The 95% CI of the LoAs for each plot is shown with a yellow shade around LoA in the plots. The BA plot statistics are summarized in Table 1. From Fig. 3, we observe that the mean bias of difference between the test-retest ROI mean values (*m*_*d*_) is close to zero for short, medium and high-*T*_2_ water fraction estimates. For all three water fraction estimations, the zero level lies comfortably inside the 95% CI of the *m*_*d*_. The test retest differences lie within the LoA and its 95% CI.

**Fig 3.**
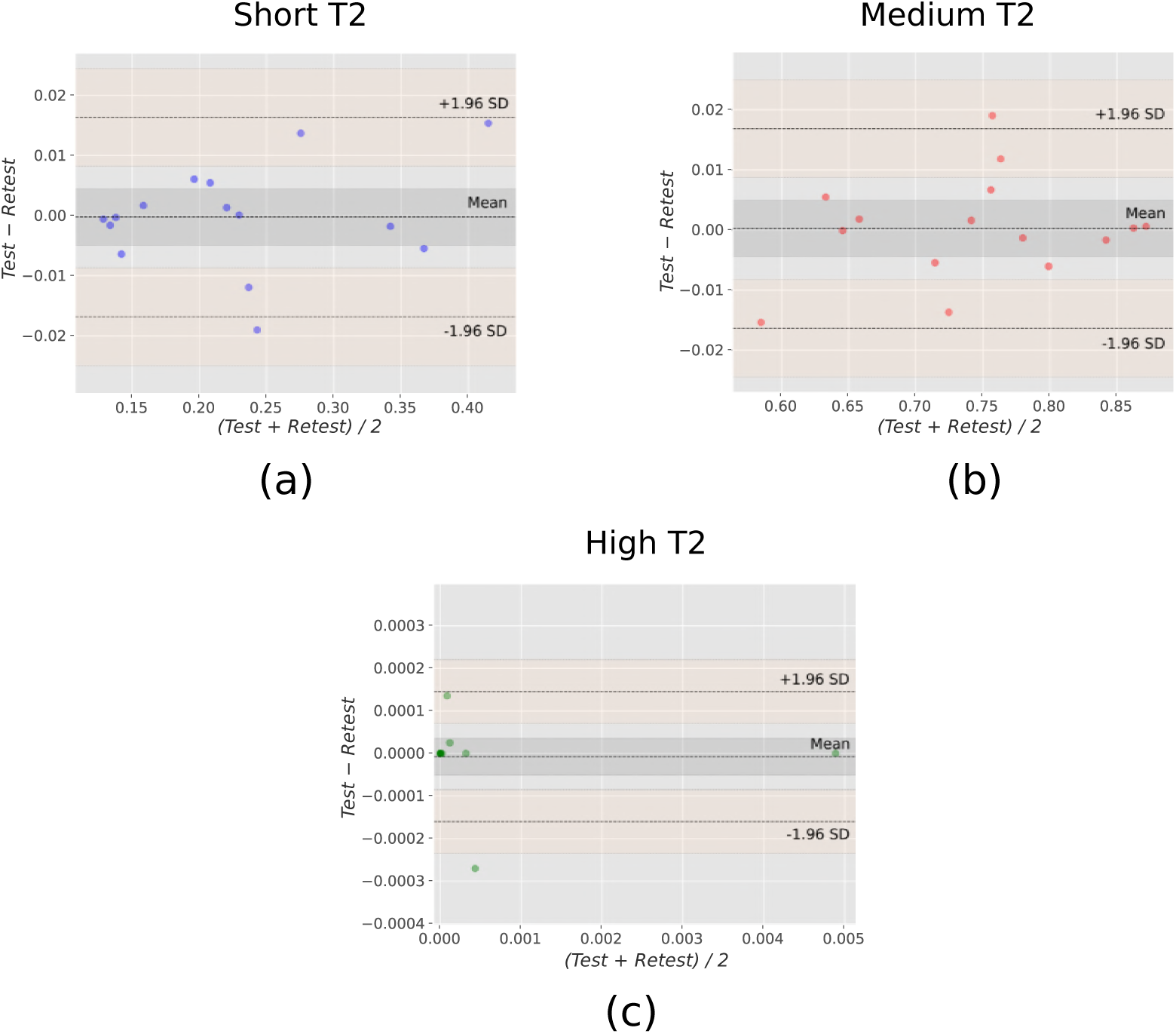
Bland-Altman plots comparing test-retest estimations. Bland-Altman (BA) plots for (a) short, (b) medium and (c) high-*T***2** water fraction estimations are shown here. 15 ROIs (see Fig. 1) are evaluated for repeatability of the estimated water fractions in 4 healthy controls. The mean level (*m*_*d*_) is the mean of the differences between test and retest values. The gray region around *m*_*d*_ is its 95% confidence interval (CI). The limits of agreement (LoA) levels are computed as *m*_*d*_ ± 1.96SD, where SD is the standard deviation of the difference in test and retest values. The yellow region around the LoAs are its 95% CI. The BA plot statistics are summarized in Table 1.

**Table 1.**
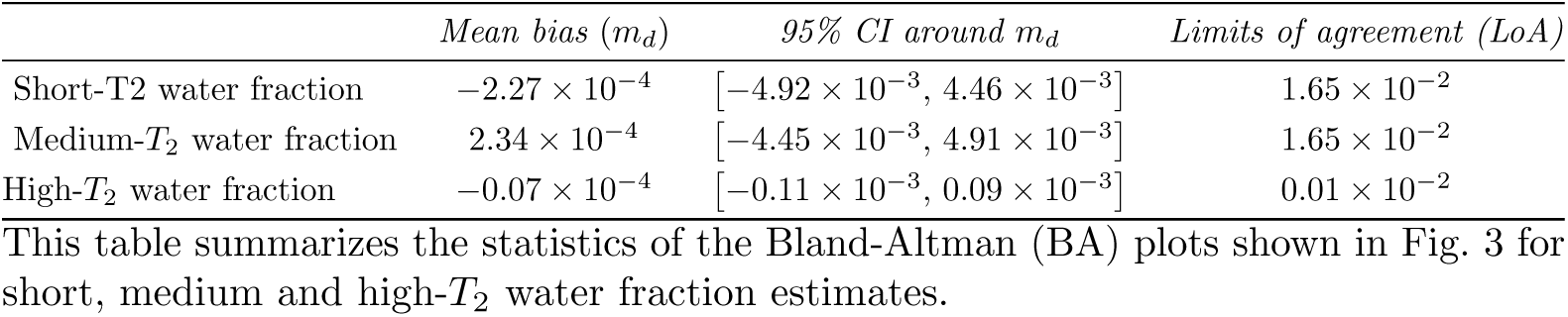
Bland-Altman plots statistics for repeatability experiment

### 3.2 Experiment-2: Application to Multiple Sclerosis (MS)

In this experiment we observed and compared the evolution of water fraction maps of the three compartments between *E*+ and *L*– MS lesion regions in 10 patients over a period of 3 years. We observed the water fraction values for *E*+ and *L*– at each scan point. In addition to it, we also observed the change in the water fraction values (for *E*+ and *L*–) between consecutive scans which is computed as: Δ*w*_*f*,*i*_ = (*w*_*f*_*sca*__*i*+1__ – *w*_*f*_*sca*__*i*__). Hence positive values indicate a gain in the water fraction values between consecutive scans. The *E*+ and *L*– group difference analysis was performed using Mann-Whitney U test.

An example illustrating the comparison between the water fraction maps for a healthy control and MS patient is shown in Fig. 4. Lesion-1 in the MS patient has a large active core, whereas a very small region of lesion-2 is active. Both lesions show indications of extensive demyelination. The medium-*T*_2_ and high-*T*_2_ water fraction maps show varying trends among the lesions, and also between the regions of the lesion undergoing active blood brain barrier breakdown and otherwise.

**Fig 4.**
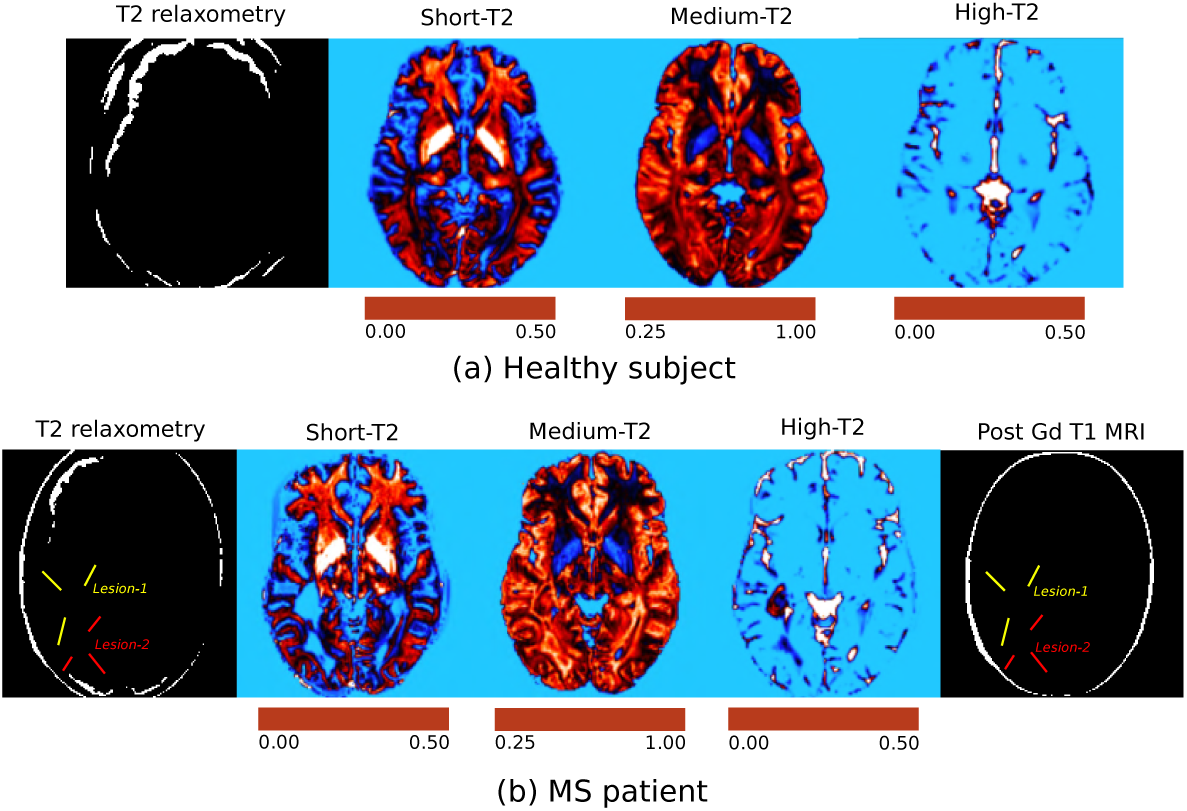
Healthy subject and MS patient. A comparison between water fraction maps for (a) healthy subject and (b) MS patient is shown.

#### Short *T*_2_ water fraction

Results for short-*T*_2_ water fraction (*w*_*s*_) maps is shown in Fig. 5. The *w*_*s*_ values of the *E*+ and *L*– lesions at all time points is shown in Fig. 5(a). The *L*– lesion regions are significantly associated with higher *w*_*s*_ values as compared to *E*+ at M00 (*p* = 0.014). However, the *w*_*s*_ distribution of *E*+ and *L*– regions at the end of 3 years are similar. The results of the change in *w*_*s*_ values (Δ*w*_*s*_) between consecutive scans in shown in Fig. 5(b). Largely negative Δ*w*_*s*,0_ values for *E*+ lesion regions suggests increased *w*_*s*_ values between M00 and M03. After M06 we observed less changes in *w*_*s*_ values in *E*+ lesion regions. The observed change in *w*_*s*_ values for the *L*– lesions regions was very less throughout the 3 year period. Only A*w*_*s* 0_ values *E*+ and *L*– were significantly different (*p* = 0.009).

**Fig 5.**
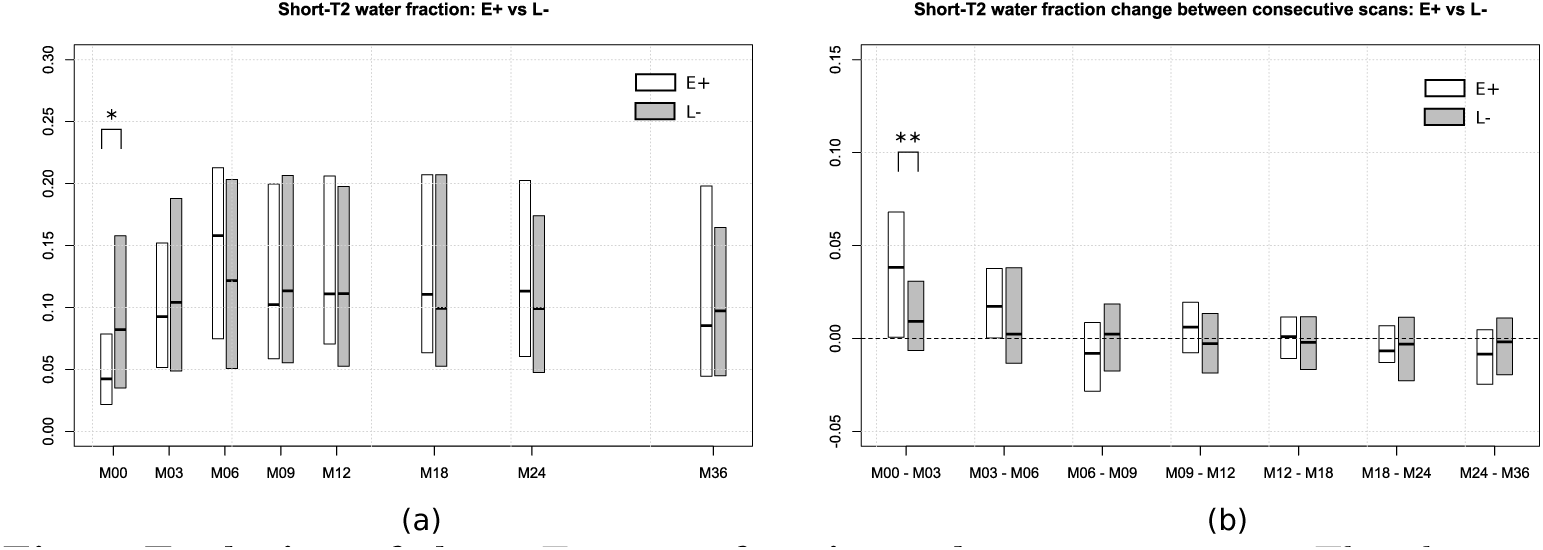
Evolution of short *T*_2_ water fraction values over 3 years. The plots in this figure show the median and upper and lower quartile levels of the data. The short-*T*_2_ water fraction value (*w*_*s*_) at each scan is shown in Fig. 5(a). The change in *w*_*s*_ between consecutive scans is shown in Fig. 5(b). Significant group differences between groups are shown using * (*p* < 0.05) and ** (*p* < 0.01).

#### Medium-*T*_2_ water fraction

The evolution of the medium-*T*_2_ water fraction (*w*_*m*_) in *E*+ and *L*– lesion regions is shown in Fig. 6(a). Although the w_m_ values for both groups reduce slightly at the end of 3 years, there is no evidence of difference between *E*+ and *L*– with respect to the change in *w*_*m*_ values between successive scans (refer Fig. 6(b)).

**Fig 6.**
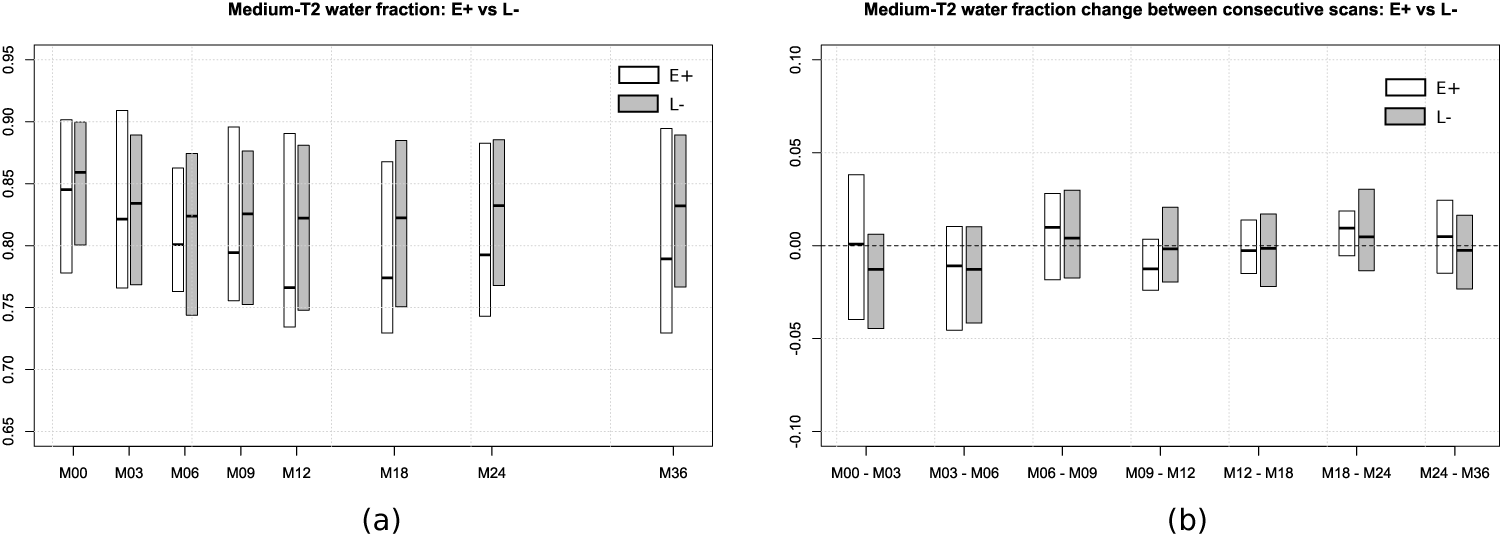
Evolution of medium *T*_2_ water fraction values over 3 years. The plots in this figure show the median and upper and lower quartile levels of the data. The medium-*T*_2_ water fraction value (*w*_*m*_) at each scan is shown in Fig. 6(a). The change ir *w*_*m*_ between consecutive scans is shown in Fig. 6(b). Significant group differences between groups are shown using * (*p* < 0.05) and ** (*p* < 0.01).

#### High-*T*_2_ water fraction

The high-*T*_2_ water fraction (*w*_*h*_) values for *E*+ and *L*– lesion regions are shown in Fig. 7(a). The *E*+ lesion regions are significantly associated with a higher value of *w*_*h*_ as compared to the *L*– population at M00 (*p* = 0.002). Largely negative Δ*w*_*h*,0_ values (refer Fig. 7(b)) for *E*+ lesion regions suggest a decrease in their *w*_*h*_ values between M00 and M03. Subsequently, *E*+ lesion regions undergo negligible change between consecutive scans. *L*– lesion regions show negligible change in their *w*_*h*_ values throughout the period of the study. The change in *w*_*h*_ values were found to be significantly different between *E*+ and *L*– lesion regions for scan periods M00-M03, M03-M06 and M06-M09 with p-values of 0.008, 0.011 and 0.11 respectively.

**Fig 7.**
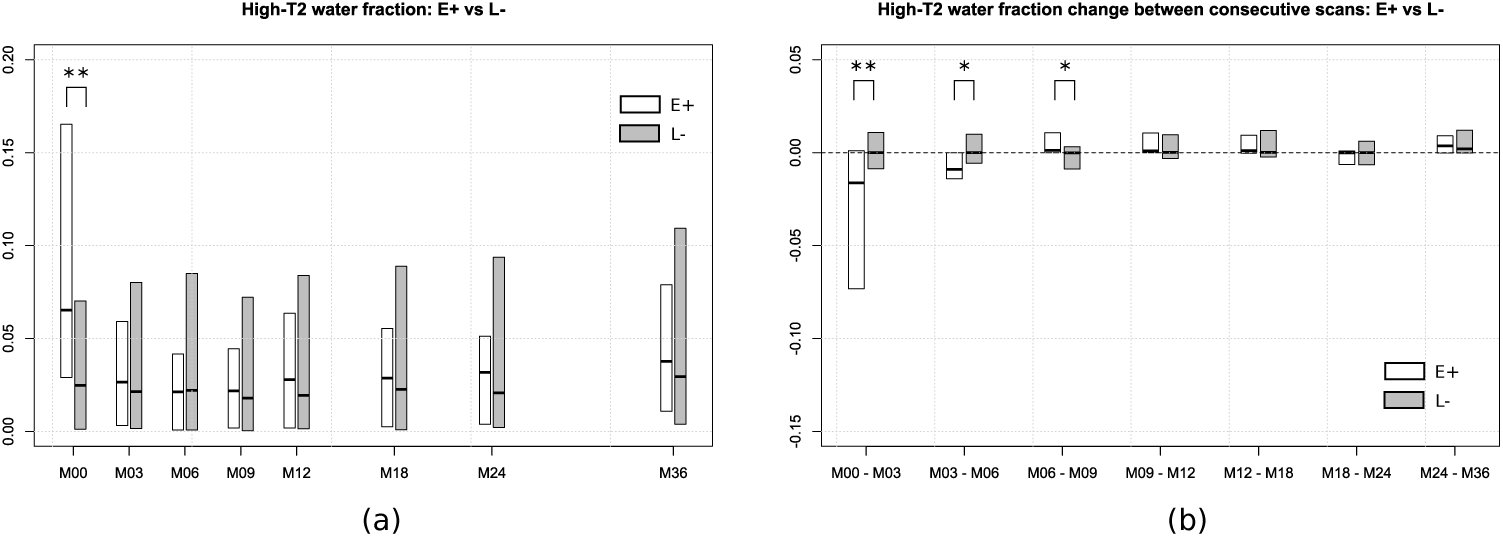
Evolution of high *T*_2_ water fraction values over 3 years. The plots in this figure show the median and upper and lower quartile levels of the data. The high-*T*_2_ water fraction value (*w*_*h*_) at each scan is shown in Fig. 7(a). The change in *w*_*h*_ between consecutive scans is shown in Fig. 7(b). Significant group differences between groups are shown using * (*p* < 0.05) and ** (*p* < 0.01).

It is also important to observe the effect size of the datasets when observing the p-values for group differences. Whereas the p-value conveys information on the strength of the water fraction values to reject the null hypothesis, the effect size is a measure of the magnitude of the difference. We used the common language (CL) effect size statistic and the associated p-values to observe the group differences [22, 23]. The CL effect size for the groups that were found to be significantly different from the Mann-Whitney U test is shown in Table 2. In this work, the CL effect size value denotes the percentage of times the wf value of *L*– is higher than *E*+ when both samples are selected at random from each group. It can hence be interpreted as the probability of superiority of *L*– over *E*+ for a measurement.

**Table 2.** Common language effect size and p-values

| Value | p-value | CL effect size |
| --- | --- | --- |
| Short-$T_2$ water fraction ($w_s$) at M00 | 0.014 | 65.05% |
| $\Delta w_s$ between M00 and M03 | 0.009 | 34.25% |
| High-$T_2$ water fraction ($w_h$) at M00 | 0.002 | 31.51% |
| $\Delta w_h$ between M00 and M03 | 0.008 | 66.08% |
| $\Delta w_h$ between M03 and M06 | 0.011 | 65.51% |
| $\Delta w_h$ between M06 and M09 | 0.011 | 34.55% |

The common language (CL) effect size are reported for the measurements which were significantly different for $E+$ and $L-$ . In this work, the CL effect size denotes the percentage of times a randomly selected sample from $L-$ is greater than a random selection from $E+$ group.

## 4 Discussion

The test-retest experiment results show that the quantitative MRI markers estimated by the proposed method is repeatable. For all the markers estimated, the zero level was comfortably inside the 95% confidence interval of the mean difference observed between the test and retest values of the marker for the 15 ROIs (refer the Bland-Altman plots in Fig. 2). Hence there are no noticeable systematic changes in the estimated markers for the test and retest data [24].

In Section 3.2 the evolution of water fraction markers in MS lesion regions which are undergoing active blood brain barrier breakdown are compared to the MS lesion regions in the later stages. At the baseline scan, the *E*+ lesion regions are prone to having lower short-T2 water fraction values as compared to *L*– regions (*p* = 0.014, CL = 65.05%). This might indicate that regions of the lesion undergoing active blood brain barrier breakdown have undergone greater demyelination. However, there seems to be no significant difference between the two groups with respect to the short-T2 water fraction values for all scans three months after the baseline. The *E*+ lesion regions tend to have significantly higher values for high-*T*_2_ water fraction as compared to *L*– at the baseline scan (*p* < 0.01, CL = 31.51%). The demyelination of MS lesions is accompanied by inflammation due to increased macrophage intervention [19]. This might explain the low and high values of short-T2 and high-*T*_2_ water fraction values observed at the baseline scan for *E*+ lesion regions. The gain in short-T2 water fraction values for *E*+ lesion regions between the scans at M00 and M03 is significantly greater than that for *L*– (p < 0.01, CL = 34.25%). The increase in short-*T*_2_ values in this period is accompanied by a considerable drop in high-*T*_2_ values for the *E*+ lesion regions. The drop in high-*T*_2_ water fraction values between the baseline and the next scan for *E*+ is significantly larger than that of *L*– (p < 0.01, CL = 66.08%). All *E*+ lesion regions incur a decrease in high-*T*_2_ values over the first six months of observation. Hence, the observations from the change in water fraction values between consecutive scans are: (i) although both groups show indications of remyelination, *E*+ lesion regions undergo significantly greater remyelination as compared to *L*– between the baseline and scan at M03, (ii) there is a considerable reduction of inflammation in *E*+ over the first three months but *L*– show little or no change in this aspect. The myelination activity of *E*+ and *L*– is similar three months after baseline. However, the inflammation activity seems to continue for 9 months from baseline scan time. Some *L*– lesion regions observed at the baseline scan would have been in the *E*+ stage at some point of time. This explains the similar water fraction values for *E*+ and *L*– by the end of three years. Unlike short-*T*_2_ and high-*T*_2_ water fraction values, the medium-*T*_2_ water fraction values for *E*+ and *L*– lesion regions never show any significant difference, which might be attributed to the fact that in terms of *T*_2_ values considered, the medium-*T*_2_ water pool is highly heterogeneous. It conveys information on unmyelinated axons, glia, interstitial and extra-cellular matters [10].

Our study on MS patients has certain limitations. First, the clinical data available was not of the highest quality possible due to acquisition time constraints in a clinical setting. *T*_2_ relaxometry data with a higher number of echoes and shorter echo times are favorable for multi-compartment models. Second, the time gap between the first and second scan was three months. A shorter interval between successive acquisitions would

## 5 Conclusion

In this work we proposed a multi-compartment *T*_2_ relaxometry model to obtain quantitative estimates on brain tissues with short, medium and high *T*_2_ relaxation times. Test retest experiment results showed that the proposed method does not suffer from any noticeable systematic changes in terms of the markers estimated. The study of the evolution of multi-compartment *T*_2_ relaxometry markers on 10 MS patients over a period of 3 years had two important observations: (*i*) the markers have the potential to distinguish between gadolinium enhanced and non-enhanced regions in MS lesions and (*ii*) both lesion regions have similar water fraction values by the end of the third year and show little distinction after 3 months from baseline scan. The later observation from the longitudinal study shows that the biomarkers obtained from this model explains the MS lesion evolution along the lines reported in the pathological and radiological studies on MS lesions [18, 19]. The former observation on the other hand motivates us to investigate methods by which biomarkers from multi-compartment models based on advanced MRI techniques can help distinguish between lesion regions undergoing active blood brain barrier breakdown without injection of contrast enhancers in MS patients, which we shall investigate in our future studies.

## Acknowledgments

The clinical MS study was supported by a grant from Fondation pour l’Aide a la Recherche sur la Sclerose en Plaques (ARSEP). The authors acknowledge Neurinfo MRI research facility for logistic support and MRI data acquisition. This research was supported in part by the grants NIH R01 EB019483 and NIH R01 NS079788.

## Footnotes

1 https://github.com/Inria-Visages/Anima-Public

